# Human chorionic gonadotropin decreases cerebral cystic encephalomalacia and parvalbumin interneuron degeneration in a pro-inflammatory model of mouse neonatal hypoxia-ischemia

**DOI:** 10.1101/2024.03.27.587006

**Authors:** Ben Miller, Alexander Crider, Bhooma Aravamuthan, Rafael Galindo

**Affiliations:** Department of Neurology, Division of Pediatric & Developmental Neurology, Washington University School of Medicine, St. Louis, MO, USA 63110

**Author notes:** Corresponding author: Rafael Galindo, MD., PhD. Department of Neurology, Box 8111 Washington University School of Medicine 660 South Euclid Avenue St. Louis, MO 63110. These authors contributed equally (Ben Miller, Alexander Crider).

**Keywords:** newborn, brain, injury, parvalbumin, interneuron, lipopolysaccharide, encephalomalacia, ischemia, hypoxia

## Abstract

The pregnancy hormone, human chorionic gonadotropin (hCG) is an immunoregulatory and neurotrophic glycoprotein of potential clinical utility in the neonate at risk for cerebral injury. Despite its well-known role in its ability to modulate the innate immune response during pregnancy, hCG has not been demonstrated to affect the pro-degenerative actions of inflammation in neonatal hypoxia-ischemia (HI). Here we utilize a neonatal mouse model of mild HI combined with intraperitoneal administration of lipopolysaccharide (LPS) to evaluate the neuroprotective actions of hCG in the setting of endotoxin-mediated systemic inflammation. Intraperitoneal treatment of hCG shortly prior to LPS injection significantly decreased tissue loss and cystic degeneration in the hippocampal and cerebral cortex in the term-equivalent neonatal mouse exposed to mild HI. Noting that parvalbumin immunoreactive interneurons have been broadly implicated in neurodevelopmental disorders, it is notable that hCG significantly improved the injury-mediated reduction of these neurons in the cerebral cortex, striatum and hippocampus. The above findings were associated with a decrease in the amount of Iba1 immunoreactive microglia in most of these brain regions. These observations implicate hCG as an agent capable of improving the neurological morbidity associated with peripheral inflammation in the neonate affected by HI. Future preclinical studies should aim at demonstrating added neuroprotective benefit by hCG in the context of therapeutic hypothermia and further exploring the mechanisms responsible for this effect. This research is likely to advance the therapeutic role of gonadotropins as a treatment for neonates with neonatal brain injury.

**Key points:**

- Intraperitoneal administration of human chorionic gonadotropin (hCG) decreases lipopolysaccharide (LPS)-augmented hypoxic-ischemic neurodegeneration in the term-equivalent mouse neonate
- Pretreatment with hCG reduces LPS-mediated cystic encephalomalacia of the cerebral cortex and ameliorates hippocampal tissue loss after neonatal hypoxia-ischemia (HI)
- hCG decreases LPS+HI-mediated parvalbumin immunoreactive interneuron loss in the cerebral cortex, hippocampus and dorsal striatum.
- hCG decreases LPS+HI-augmented microglial Iba1 immunoreactivity in the cerebral cortex and hippocampus.

## Introduction

Neonatal cerebral hypoxia-ischemia (HI) is a major cause of long-term neurodevelopmental disability in children, and it is associated with an increased risk for epilepsy, cerebral palsy and neurodevelopmental disorders. Importantly, inflammation contributes significantly to the morbidity of perinatal HI and its pro-degenerative effects are not prevented fully by therapeutic hypothermia (TH), the only available treatment in HI for affected late preterm and term neonates (Hagberg et al., 2015). For example, preclinical studies in neonatal rodents and piglets demonstrate lack of neurological benefit of TH in injured animals sensitized by lipopolysaccharide (LPS), a well-known pro-inflammatory molecule found in the outer membrane of gram-negative bacteria (Martinello et al., 2022; Osredkar et al., 2014). This evidence is supported by the absence of significant alterations in the levels of pro-inflammatory cytokines by TH relative to normothermia in LPS-sensitized rodents exposed to HI (Chevin et al., 2016). Similar results have been found in humans in which hypothermia does not confer as much benefit and may potentially be detrimental in neonates with encephalopathy affected by infection or exposed to a pro-inflammatory environment in utero (Danladi and Sabir, 2021; Wintermark et al., 2010). Therefore, neuroprotective therapies that target the negative inflammatory actions contributing to newborn brain injury may offer added therapeutic benefit in newborns affected by HI and/or exposed to pro-inflammatory conditions that further augment the degree of cerebral injury. Indeed, recent research efforts have explored treatments that target the dysregulated immune response observed in perinatal brain injury with varying levels of success, as it is the case with agents such as erythropoietin, stem cells and melatonin (Molloy et al., 2022; Pedroza-Garcia et al., 2022).

The pregnancy hormone, human chorionic gonadotropin (hCG) is a growth-promoting glycoprotein belonging to the cystine-knot growth factor superfamily with properties that make it a potentially novel therapeutic tool against the neurodegenerative contribution of inflammation to perinatal brain injury. Evidence supporting this hypothesis include hCG’s known role in modulating the immune response during pregnancy (Schumacher et al., 2014). hCG is known to reduce the phagocytic action of macrophages, decrease the activation of T-lymphocytes and inhibit B-cell antibody synthesis (Nikolaevich et al., 1991; Shirshev, 1997; Zhang et al., 2003). At the molecular level, hCG has been shown to reduce the production of pro-inflammatory cytokines such as tumor necrosis factor, IL-6 and interferon gamma and to increase the synthesis of anti-inflammatory molecules such as IL-10 and leukocyte inhibitory factor (Aghajanova, 2004; Aghajanova, 2010; Espey and Richards, 2002; Khil et al., 2007; Wan et al., 2008). Interestingly, all the above cellular processes and responses are known to contribute to the adverse neurological effects of inflammation in newborns exposed to HI. Furthermore, and aside from his anti-inflammatory actions, hCG is neuroprotective. For example, our laboratory and others have observed that hCG is able to increase the survival of neurons and glia, promote neurite outgrowth, decrease glutamate receptor-mediated excitotoxicity and protect against the degenerative effects of hypoxia-ischemia in mouse newborns and stroke in adult rats (Babahajian et al., 2019; Belayev et al., 2009; Blair et al., 2019; Movsas et al., 2017). Taken together, the immunomodulatory properties of hCG and its neurotropic actions may promote compound protection against the degenerative effects of HI potentiated by the effects of inflammation.

The potential neuroprotective effects of hCG may be global or selective for certain neuronal populations. Parvalbumin immunoreactive interneurons, particularly in the cortex, appear to be preferentially affected by acute inflammatory insults in the developing brain(Feng et al., 2021; Stolp et al., 2019; Yates et al., 2022). These neurons become hyper-excitable following systemic inflammation induced with LPS and are lost following systemic inflammation and mitigated with concomitant anti-inflammatory agent administration(Feng et al., 2021; Stolp et al., 2019; Yates et al., 2022). Furthermore, neurodevelopmental parvalbumin interneuron pathology has been implicated in the pathophysiology of neurodevelopmental disorders ranging from autism to childhood-onset dystonia(Gernert et al., 2000; Hamann et al., 2007; Juarez and Martinez Cerdeno, 2022; Song et al., 2013; Woodward and Coutellier, 2021). Therefore, it would be valuable to elucidate druggable mechanisms specifically targeted to the rescue of parvalbumin immunoreactive interneurons.

This report explores the neuroprotective actions of hCG, including on parvalbumin immunoreactive interneurons, in the setting of systemic inflammation using a mouse model of LPS-sensitized neonatal brain injury. We find that pretreatment with hCG significantly reduces hippocampal and cortical cystic neurodegeneration. This effect is associated with a reduction in the extent of microglial Iba1 immunoreactivity in these brain regions. Furthermore, we observe an improvement in the loss of parvalbumin immunoreactive neurons in cerebral cortex, dorsal striatum and CA3/dentate-gyrus region of the mouse hippocampus. These findings implicate hCG as a neuroprotective agent capable of reducing the deleterious effects of systemic inflammation in the setting of cerebral injury. Future studies should aim at examining the immunomodulatory mechanisms responsible for this effect in the context of preterm and term neonatal brain injury.

## Methods

### Animals

The care and use of all mice were in strict accordance with the National Institute of Health Guidelines on the Care and Use of Laboratory Animals. All procedures were performed according to an approved protocol by the Washington University Institutional Animal Care and Use Committee. Male and Female mice of C57BL/6J background were used. Every effort was made to segregate mice between groups such that weight and gender were of equal or nearly equal proportions at the start of the experiment. To avoid maternal confounders, equal number of pups from the control and experimental group were represented within a given litter. Control and experimental mice were injected or underwent surgery on the same day and were exposed to the hypoxia chamber at the same time. Pup health, weight and maternal care were routinely surveyed following drug exposure. Animals that lost equal or greater than 20% of their body weight were not processed for further evaluation and were euthanized.

### Lipopolysaccharide-sensitized Model of Neonatal Hypoxia-Ischemia

Postnatal day 8 mice received an intraperitoneal (IP) injection 0.6 µg/g of lipopolysaccharide (LPS) from Escherichia coli O55:B5 (Sigma-Aldrich, St. Louis MO, Catalog no. L2880) diluted in sterile 0.9% sodium chloride solution (normal saline). All IP injections were performed using a sterilized Microliter™ Hamilton syringe with 30 GA 0.5’’ PT4 style needles (Hamilton Co., Reno, NV, Catalog no. 702 RN). To evaluate the effects of LPS in our mouse model of hypoxia-ischemia (HI), control pups received an equal liquid volume per gram of weight of normal saline solution to that of the LPS-treated group (Figure 1A). Approximately 18 hours after injection, pups were anesthetized with isoflurane (4% induction, 3% maintenance) and the left internal carotid artery exposed through a neck incision. Flow through the left carotid artery was interrupted by cauterization and the wound closed with tissue glue (GLUture, Zoetis Inc, Kalamazoo, MI). Pups were then returned to the dam for three hours to recover from carotid surgery. Following the post-op recovery periods, animals were placed in a thermoregulated hypoxia chamber and exposed for 20 minutes to high grade humidified 8% oxygen (±0.02%) balanced with nitrogen at 36.5 °C. Following hypoxia exposure, pups were returned to the dam until the day they were euthanized at postnatal day 16.

**Figure 1.**
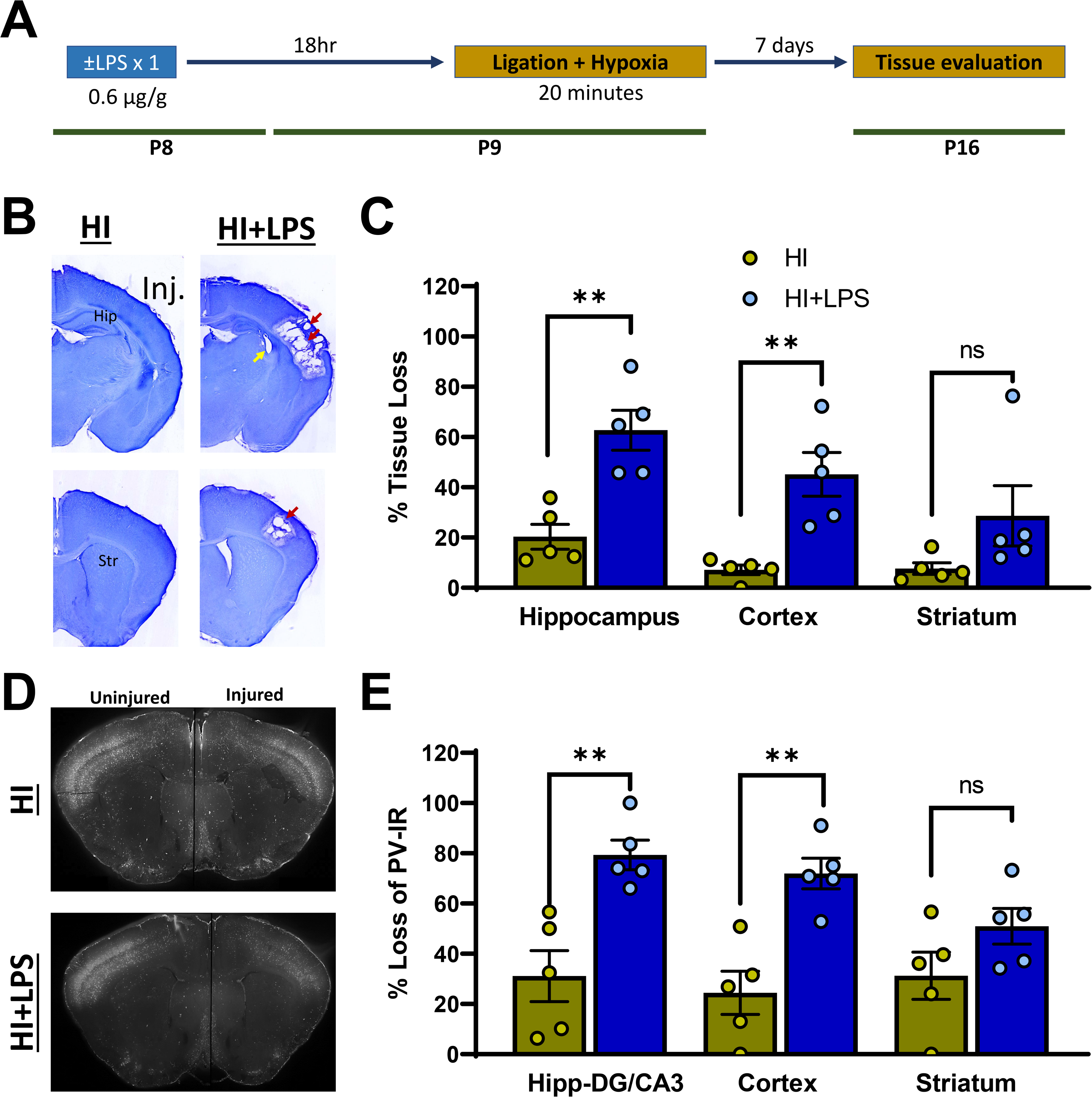
LPS sensitizes the neonatal brain to the effects of HI. **A.** LPS-augmented neonatal brain injury paradigm. **B.** Representative cresyl violet stained sections of the ipsilateral injured (Inj.) brain at the level of the dorsal hippocampus (Hip; top) and dorsal striatum (Str; bottom) from HI-only and HI+LPS treated mice. Note the presence of cystic cavitation adjacent to the hippocampus (yellow arrow) and associated cerebral cortex (red arrows) seen predominantly in the LPS-treated mice. **C.** Quantitative comparison of the amount of tissue loss between HI-only and HI+LPS treated animals in the hippocampus, cortex and striatum. Percent tissue loss was calculated by dividing the area of the injured region over the uninjured side. **D.** Parvalbumin immunoreactivity (PV-IR) in representative sections of the dorsal striatum. Note the significant decrease in PV-IR in the ipsilateral injured cortex of LPS-treated mice (yellow arrowheads). **E.** Quantitative evaluation of the percent loss of PV-IR in the dentate gyrus/CA3 region (Hipp-DG/CA3), para-striatal cortex and dorsal striatum relative to the uninjured hemisphere. **p < .01 by unpaired two-tailed Student’s t-test (ns; non-significant).

### hCG and Vehicle Injections

To examine the neuroprotective effects of hCG in a pro-inflammatory model of neonatal HI, we used our LPS-sensitized model of HI described above and delivered IP 1.5 IU/g of hCG diluted in normal saline (Sigma-Aldrich, St. Louis, MO, Catalog no. CG-10) or normal saline alone as vehicle control two hours prior to LPS administration (Figure 2A). Equal liquid volumes per gram of pup weight to that of hCG group were given to the control animals. hCG-treated and control-treated animals were injected and received carotid ligation in a sequential manner to avoid potential confounders related to timing of administration. Similarly, both groups were exposed simultaneously to hypoxia and segregated in proportional numbers and gender to the assigned dam to prevent maternal care confounders. Animal weight was obtained daily during the first 3 days from the start of the experiment and then at 6 and 8 days post drug treatment. Animals were euthanized on postnatal day 16 and processed for histology.

**Figure 2.**
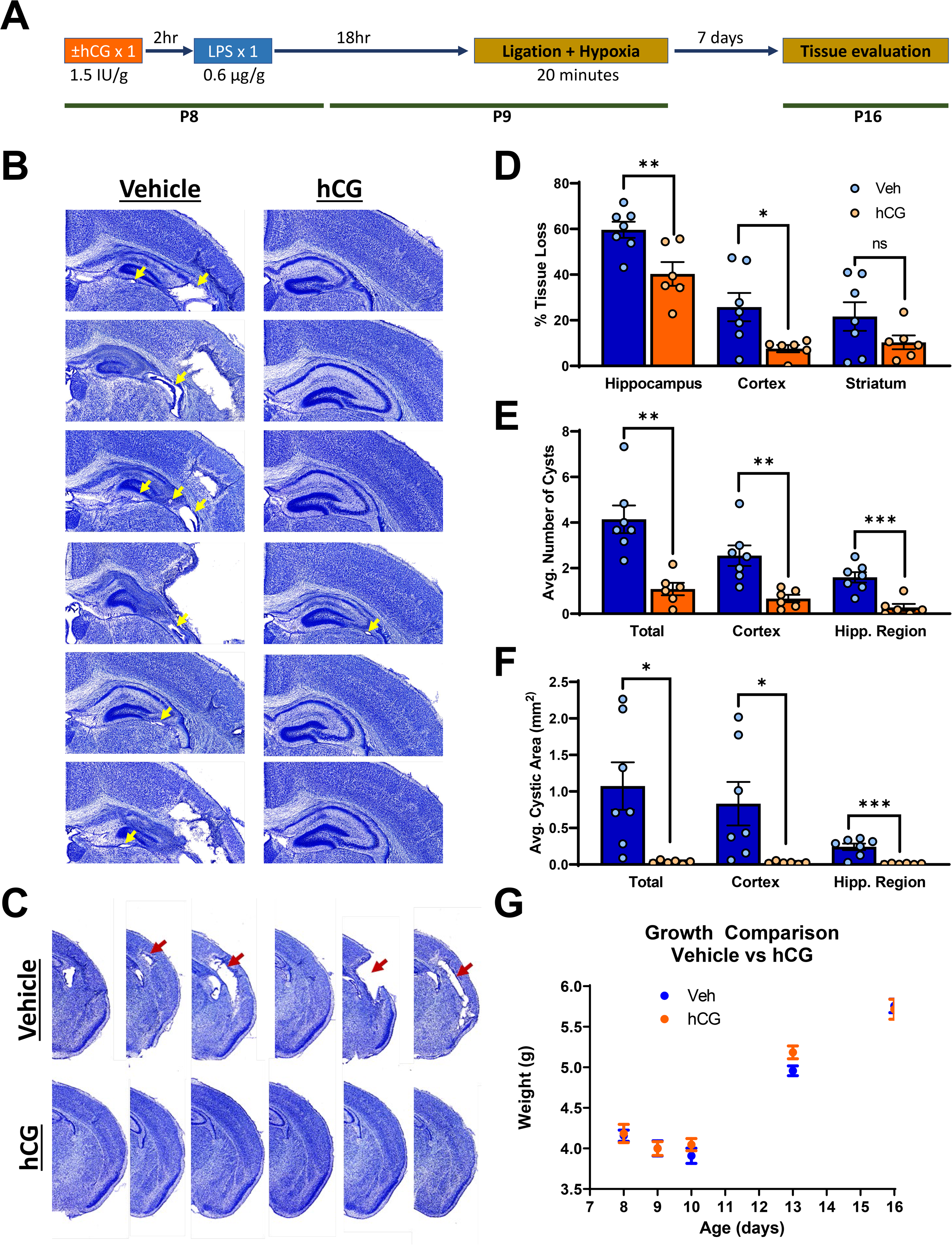
hCG protects against LPS-augmented hippocampal and cortical degeneration following HI. **A.** Experimental paradigm for hCG prophylaxis of LPS-augmented brain injury. **B.** Cresyl violet stained sections of hippocampus from hCG treated animals and controls injected with normal saline vehicle. Note severe cystic degeneration in control mice (yellow arrows). **C.** Cresyl violet stained sections demonstrating cystic degeneration (red arrows) in the cortex of control mice not seen in hCG treated subjects. **D.** Quantitative comparison of the amount of tissue loss between hCG treated and vehicle treated control animals in the hippocampus, cortex and striatum. Percent tissue loss was calculated by dividing the area of the injured region over the uninjured side. **E.** Comparison of the absolute number of cystic lesions between hCG treated subjects and vehicle treated controls. **F.** Comparison of the total area of the cystic regions between hCG treated subjects and vehicle treated controls. **G.** Weight of the mice over the course of the experiment. *p < .05, **p < .01, ***p < .001 by unpaired two-tailed Student’s t-test.

### Histology and Tissue Injury Assessment

Postnatal day 16 hCG- and vehicle-treated pups were deeply anesthetized with an IP injection of sodium pentobarbital (Fatal-Plus; 200 mg/kg) and perfused transcardially with ice-cold phosphate buffer saline (PBS) containing 3 U/mL of heparin. Brains were then removed and fixed by immersion in 4% paraformaldehyde diluted in PBS for 48 hours followed by cryoprotection with 30% sucrose in PBS. Coronal brain slices of 50 µm thickness were obtained using a slicing freezing microtome, stained with cresyl violet and scanned using a Hamamatsu NanoZoomer 2.0 digital slide scanner (Hamamatsu Photonics K.K., Shizuoka, Japan). Six coronal sections per quantified area (spaced 50 µm apart) were used for each animal to examine the extend of cerebral injury. Areas evaluated were the dorsal striatum at level of the genu of the corpus callosum, cerebral cortex and the dorsal hippocampus. Cortical injury was quantified using NDP.view2 software (Hamamatsu Photonics K.K., Shizuoka, Japan) on the sections containing the dorsal hippocampus unless otherwise specified. Using a unique animal number identifier, the investigators were blinded to the treatment condition following hypoxia and throughout the quantification procedure. Extent of tissue loss was determined by calculating the percent area difference in the injured, ischemic (left), hemisphere to the uninjured, non-ischemic, hemisphere as previously described (Movsas et al., 2017). A cystic lesion was defined as a well circumscribed area devoid of neural tissue.

### Immunohistochemical staining and quantification

To evaluate the injury-mediated effects of parvalbumin GABAergic interneurons and Iba1 positive microglia, we utilized a free-floating method of immunohistochemical staining. Primary antibodies used were mouse anti-parvalbumin (1:500; MAB1572, MilliporeSigma, Burlington, MA) and rabbit anti-Iba1 (1:500; #019-1974, FUJIFILM Wako, Richmond, VA). Coronal 50 µm sections were washed in PBS, blocked with goat serum in permeabilizing buffer (10% normal goat serum with 0.5% Triton X-100 in PBS) for 2 hours followed by overnight immersion with primary antibody in incubation solution (5% goat serum with 0.3% Triton X-100 in PBS) at 4°C. The following day, tissue was washed in PBS and immersed in goat anti-mouse or anti-rabbit IgG (H+L) cross-adsorbed Alexa Fluor™ 568 (1:500; ThermoFisher Scientific, Waltham, MA) prepared in incubation solution for 2 hours at room temperature. Brain sections were then mounted in Fluoromount (ThermoFisher Scientific, Waltham, MA) and scanned using a Hamamatsu NanoZoomer 2.0 digital slide scanner (Hamamatsu Photonics K.K., Shizuoka, Japan).

Images were calibrated for identical brightness levels post-imaging using NDP.view2 software (Hamamatsu Photonics K.K., Shizuoka, Japan). This brightness allowed for optimal viewing of neuronal staining without contamination from background staining across all brain slices. Images were further analyzed using ImageJ. Regions of interest (ROIs) were drawn bilaterally (injured and non-injured side). These ROIs encompassed the primary motor and primary sensory cortices using the corpus callosum as a landmark. The same ROIs were used on to quantify parvalbumin immunoreactivity on every slice assessed to ensure comparable areas were assessed for each brain. Slices were set to 8-bit grayscale and then thresholded identically. ROIs were then overlayed on these thresholded slices. For quantifying parvalbumin immunoreactivity in the sensorimotor cortex, we calculated the percent area occupied by the thresholded signal within each sensorimotor cortex ROI. For quantifying parvalbumin immunoreactivity in the striatum and dorsal hippocampus, individual neuron bodies were manually counted for each ROI for each slice. Notably, throughout all quantification, the experimenter was blinded to the experimental condition.

### Statistical methods

Statistical analysis was performed with GraphPad 9 Prism software. Data is calculated and presented as mean ± standard error of the mean. Two-tailed Student’s t-test was used for two-group comparisons. Statistical significance was set at .05, where *p < .05, **p < .01, ***p < .001.

## Results

### LPS sensitizes the neonatal brain to the effects of HI

To evaluate the prodegenerative effects of LPS in neonatal hypoxia-ischemia (HI), postnatal day 8 mice underwent a single intraperitoneal (IP) administration of 0.6 µg/g of LPS (N=5) or normal saline control solution (N=5). The next day, all mice underwent unilateral carotid ligation followed by 20 minutes of 8% FiO2 hypoxia and their brains were examined 7 days post HI (Fig. 1A). HI mice treated with LPS demonstrated obvious increases in tissue injury relative to HI alone as noted by the presence of para-hippocampal and cortical cystic degeneration (Fig. 1B). Quantitatively, HI-LPS mice had significant increases in hippocampal and cortical tissue loss relative to the animals only exposed to HI (Fig. 1C). Immunohistochemical evaluation of parvalbumin (PV) interneurons revealed that HI-LPS mice had significantly decreased PV immunoreactivity (IR) compared to mice exposed to HI alone, particularly in the cerebral cortex and dentate gyrus/CA3 region of the hippocampus in the ischemic hemisphere ipsilateral to carotid ligation (Fig. 1D-E).

### Intraperitoneal administration of hCG protects against LPS-augmented hippocampal and cortical degeneration following HI

We next evaluated the antidegenerative effect of hCG treatment in the HI-LPS paradigm by injecting P8 mice with vehicle control solution (n=8; 5 females and 3 males) or 1.5 IU/g of hCG two hours prior to LPS administration (n=7; 5 females and 2 males; Figure 2A). One animal from each group died either during the hypoxia period or shortly after. Qualitatively, hCG-treated mice showed a relative preservation of hippocampal cytoarchitecture and an apparent decrease in para-hippocampal and cortical cystic degeneration compared to vehicle control-treated mice (Fig. 2B-C). Quantitative assessment of tissue loss demonstrated an hCG-mediated reduction in the degree of hippocampal and cortical but not striatal injury (Fig. 2D). Similarly, the number and average size of cystic lesions in the cerebral cortex and hippocampal region was reduced by hCG (Fig. 2E-F). Despite this neuroprotective effect, the overall growth of hCG-treated mice did not differ from mice treated with control solution (Fig. 2G).

### hCG ameliorates the reduction in parvalbumin-positive immunoreactivity in the cortex, striatum and hippocampus following LPS-augmented HI

Given the notable observed decrease in PV-IR of the ischemic hemisphere in the HI-LPS paradigm, we studied the effect of hCG in this neuronal population. Vehicle-treated mice again demonstrated a notable reduction in PV-IR in the ischemic cerebral cortex and striatum relative to the contralateral hemisphere (Fig. 3A-B). hCG treatment produced a significant relative reduction in the loss of PV-IR in both brain regions in the ischemic hemisphere (Fig. 3A-B). Examination of the number of PV+ neurons in the hippocampus also revealed an hCG-mediated improvement in the loss of PV-IR specifically in the dentate gyrus and CA3 region of the hippocampus (Fig. 3 C-D).

**Figure 3.**
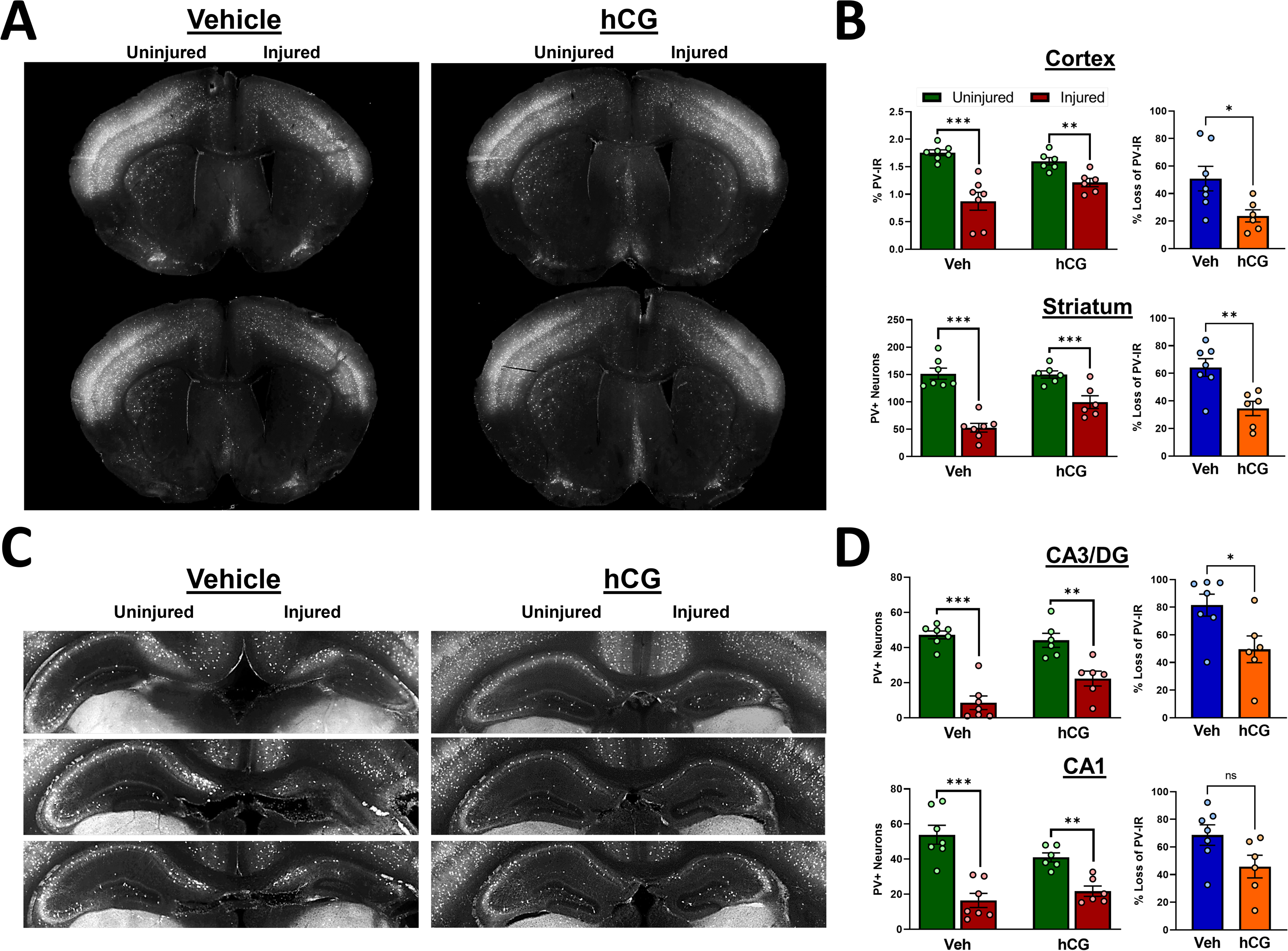
hCG preserves parvalbumin immunoreactivity in mice subjected to LPS-augmented HI. **A.** Parvalbumin immunohistochemistry of representative striatal sections of vehicle treated controls and hCG treated subjects. Note preserved parvalbumin staining on the injured (left) side of the hCG treated animal. **B.** Quantitative comparison of the percentage of parvalbumin staining between in the cortex of vehicle treated controls and hCG treated subjects (top). Number of parvalbumin positive neurons in the striatum of vehicle treated controls and hCG treated subjects. **C.** Parvalbumin immunohistochemistry of representative dorsal hippocampus sections of vehicle treated controls and hCG treated subjects. Not preserved number of parvalbumin positive neurons in the injured (left) side of the hCG treated mice. **D.** Quantitative comparison of the number of parvalbumin positive neurons in the CA3/Dentate gyrus (DG) region of the hippocampus (top) and in the CA1 region of the hippocampus (bottom). *p < .05, **p < .01, ***p < .001 by unpaired two-tailed Student’s t-test.

### hCG decreases hippocampal Iba-1 immunoreactivity after LPS-augmented HI

Lastly, we examined the effect of hCG on Iba1+ microglia following combined LPS-HI. Iba1-IR was particularly abundant in the ischemic hippocampus (Fig. 4A). Treatment with hCG demonstrated a relative decrease in the amount of Iba1-IR compared to HI-LPS mice treated with control vehicle solution (Fig. 4A-B). A similar effect was observed in the cerebral cortex primarily in the region surrounding the cystic lesions.

**Figure 4.**
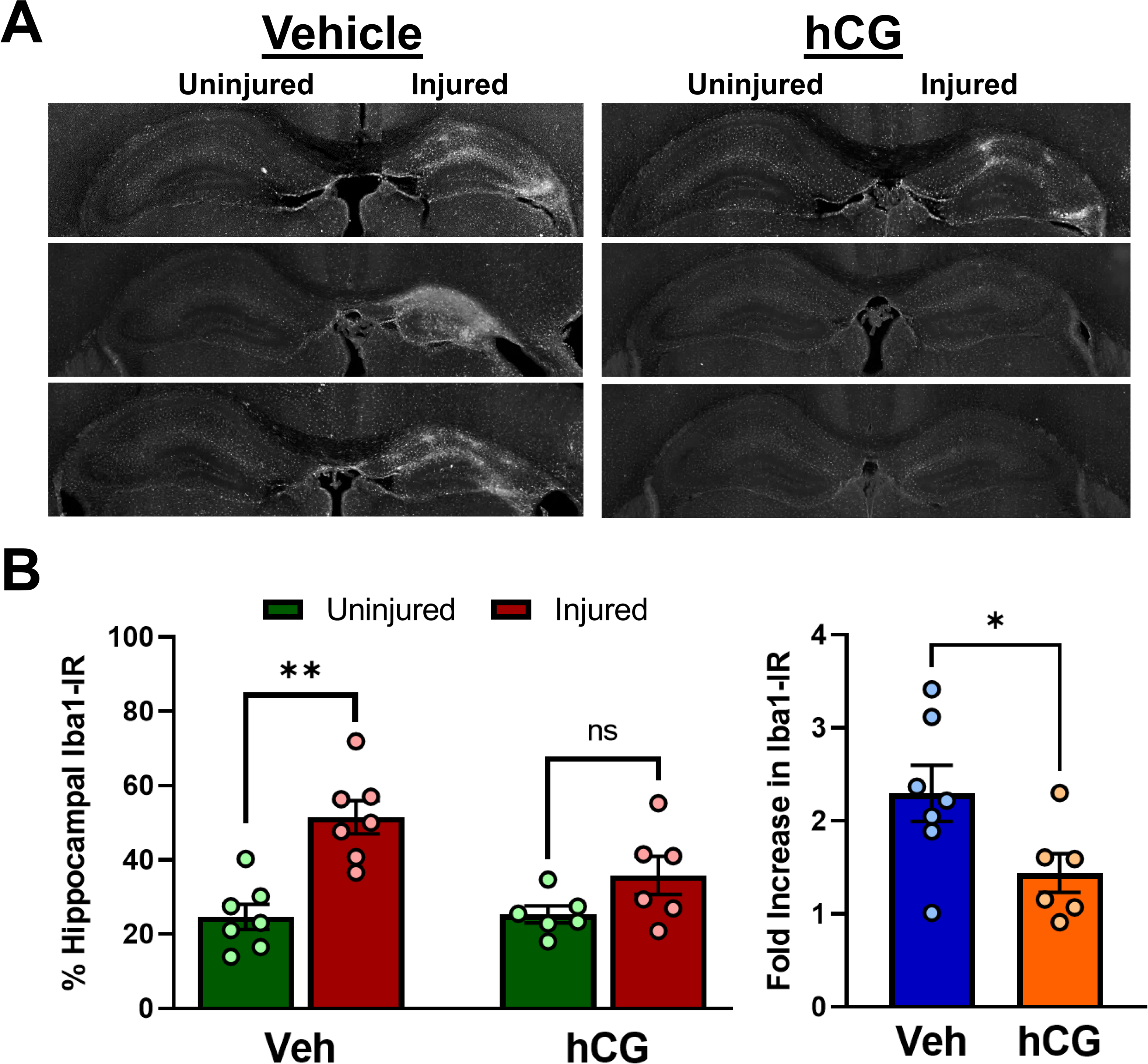
hCG decreases hippocampal Iba-1 immunoreactivity after LPS-augmented HI. **A.** Iba-1 immunohistochemistry of representative dorsal hippocampal sections of vehicle treated controls and hCG treated subjects. Note the decreased Iba-1 staining in the injured (left) side of the hCG treated mice. **B.** Quantitative comparison of the amount of Iba-1 immunoreactivity between vehicle treated controls and hCG treated animals. *p < .05, **p < .01 by unpaired two-tailed Student’s t-test.

## Discussion

In this report we find that intraperitoneal administration of hCG shortly before the induction of a systemic inflammatory response by LPS significantly reduced the degree of cerebral degeneration from HI in the term-equivalent newborn mouse brain. This effect was further associated with a decrease in Iba1+ microglia in the injured cortex and hippocampus. Furthermore, we also observed an improvement in the injury-mediated loss of parvalbumin immunoreactivity in the hippocampus, striatum, and cerebral cortex demonstrating this hormone’s ability to protect specific neuronal phenotypes.

The above results indicate that hCG can decrease the neurodegenerative effects of systemic inflammation in mice exposed to cerebral hypoxia-ischemia. To our knowledge, this is the first report demonstrating the ability of a gonadotropin to ameliorate the neurodegenerative effects of inflammation in the setting of neonatal cerebral injury. hCG is a hormone highly implicated in the immune-endocrine cross talk during pregnancy and with diverse immunological actions. The elevation of hCG early in pregnancy is important in facilitating the implantation and maintenance of the embryo and later regulate the immune response to avoid the rejection of the semiallogeneic fetus (Schumacher and Zenclussen, 2019). Interestingly, the hCG-mediated immunomodulatory mechanisms involved in preserving fetal tolerance share a very similar profile to those observed in perinatal hypoxic-ischemic brain injury. Neonatal HI results in the activation of multiple early and late inflammatory cellular processes facilitated by the compromised blood brain barrier and the death of neurons and glia.

Neurodegeneration from HI triggers the activation of microglia, penetrating monocytes, lymphocytes and neutrophils and the subsequent release of pro-inflammatory cytokines producing secondary cell death (Li et al., 2017). LPS augments these processes by activating toll-like receptor 4 (TLR4) signaling in these immune elements and increasing further the peripheral and central inflammatory response (Wang et al., 2006).

Interestingly, hCG is known to favorably affect all these cellular processes. For example, hCG can enhance the death and inhibit the proliferation of neutrophils and lymphocytes (Shirshev and Kuklina, 2001; Shirshev et al., 2003). It similarly modulates and dampens the pro-inflammatory phenotype of monocytes and macrophages while decreasing the release of pro-inflammatory cytokines such as interferon-γ and TNF-α and augmenting the secretion of anti-inflammatory molecules including TNF-β, IL-10 and leukocyte inhibitory factor (Furcron et al., 2016; Khil et al., 2007; Wan et al., 2008; Yu et al., 2015). Furthermore, preclinical studies indicate that hCG has relative affinity towards modulating the pro-inflammatory effects of LPS. hCG exposure of mouse bone marrow-derived antigen presenting dendritic cells or splenocytes reduces LPS-mediated TNF-α secretion and T cell proliferation (Khil et al., 2007; Wan et al., 2008). Moreover, hCG-mediated decreases in human monocyte phagocytic activity are inhibited by TLR4 receptor blockade *in vitro* thus implicating directly hCG receptors in TLR4-mediated immune cell function (Shirshev and Zamorina, 2006). Indeed, hCG receptors are expressed in macrophages including microglia further supporting the potential direct immunomodulatory actions of gonadotropins in the injured brain (Bukovsky et al., 2003; Zhang et al., 2003). It is important to note, however, that hCG is a pleiotropic hormone capable of increasing the production of estrogen and progesterone, two sex hormones that are known to also participate in the regulation of the immune response (Schumacher et al., 2014). Therefore, although is tempting to conclude that the observed reduction in LPS-mediated cerebral injury after neonatal HI is attributed directly to the central and/or peripheral immunomodulatory and neurotrophic actions of hCG, multiple other mechanisms could be involved including indirect effects by other reproductive hormones. Future studies aimed at examining the peripheral and central inflammatory changes triggered by LPS in hCG-receptor deficient mice and neurons may help clarify the specific mechanisms involved. Our laboratory is actively testing molecular tools such as viral-mediated RNA interference to modulate hCG receptor expression which could be used to further elucidate how the activation of these receptors by gonadotropins affects the health of neurons and glia in response to perinatal ischemia and inflammation.

In addition to the observed general hCG-mediated reduction in tissue degeneration triggered by HI-LPS, we observed that this type of injury significantly decreased the amount of parvalbumin immunoreactive interneurons in the cerebral cortex, striatum and hippocampus. Importantly, this effect was ameliorated by hCG treatment in these brain regions. Parvalbumin is a calcium binding protein found in a subgroup of inhibitory interneurons. PV interneurons are responsible for the generation of high energy-demanding gamma oscillations, and they receive the greatest excitatory input of any inhibitory interneuron in the brain (Ruden et al., 2021). PV neurons play a significant role in synaptic plasticity and multiple lines of evidence indicate that they are highly susceptible to the effects of cerebral injury and inflammation (Andersen, 2022; Ruden et al., 2021). For example, significant decreases in the amount of PV-IR neurons have been observed in brains exposed to LPS during the prenatal, neonatal and adult periods (Basta-Kaim et al., 2015; Chen et al., 2022; Crapser et al., 2020; Jenkins et al., 2009; Wischhof et al., 2015). In agreement with our observations, the loss of PV-IR by LPS is found associated with microglial activation and was particularly prominent in the cerebral cortex, dentate and CA3 regions of the hippocampus (Chen et al., 2022; Jenkins et al., 2009; Wischhof et al., 2015). These changes are known to affect PV neuronal integrity and contribute to an impairment in affect and/or cognitive function (Jenkins et al., 2009; Wischhof et al., 2015). Various other studies utilizing different animal models support the notion that PV levels or PV interneuron integrity is adversely affected by increases in peripheral or central inflammation (Andersen, 2022; Ji et al., 2015; Wang et al., 2021; Zinsmaier et al., 2020). Furthermore, PV-IR neurons are decreased in the striatum and cortex of developing animals exposed to hypoxia or hyperoxia (Mallard et al., 1995; Schiavone et al., 2017). Striatal PV-IR changes are particularly interesting in the context of the motor abnormalities, like dystonia, that are commonly associated with cerebral palsy(Rice et al., 2017). Decreased PV-IR has been observed in multiple genetic mouse models of early-onset dystonia(Gernert et al., 2000; Hamann et al., 2007; Song et al., 2013). Whether a similar mechanism can lead to dystonic motor abnormalities following HI-LPS remains to be assessed.

There are multiple ways via which hCG may have mitigated PV-IR loss following HI-LPS. One is that PV interneurons were lost following HI-LPS and that hCG mitigated this neuronal loss. The other possibility is that the neurons remained intact but that parvalbumin expression within these neurons decreased following HI-LPS and that this decreased expression was amelioriated with hCG. Given the role of parvalbumin as a calcium buffer, its expression is closely tied to neuronal activity(Filice et al., 2016).

Downregulation of parvalbumin expression is thought to be associated with decreased PV interneuron activity during development, though this relationship is complex and remains under study(Caillard et al., 2000; Patz et al., 2004). To further investigate the mechanism underlying the PV-IR changes observed here, future studies could assess changes in PV interneuron activity in HI-LPS mice with and without concomitant hCG administration.

Taken together, the above evidence suggests that hCG favorably impacts PV-IR by indirectly or directly decreasing the negative effects of inflammation triggered by LPS and HI. Importantly, impairment in the integrity and/or function of these neurons has been implicated in several disorders commonly observed in individuals affected by neonatal HI including cognitive impairment, epilepsy, developmental and mood disorders (Chen et al., 2022; Fawcett et al., 2019; Filice et al., 2020; Schwaller et al., 2004). Therefore, the neuroprotective effect observed by hCG on this neuronal population has likely important clinical implications for these diseases. Future behavioral and functional studies evaluating the significance of this finding may help clarify its neurological impact.

Despite the favorable cerebral effects observed by pretreatment of hCG on LPS-sensitized neonatal HI. It is not yet known whether the neuroprotective effects of hCG persist if the hormone is administered after the induction of HI-LPS injury. Examining whether hCG remains neuroprotective in this context will broaden support for its research translation. Yet, prophylactic use of hCG is likely to have translatable clinical utility since recurrent pro-inflammatory complications from infection and other postnatal adverse events are commonly encountered in the critically ill neonate and contribute significantly to long term neurological morbidity (Berger et al., 2012; Humberg et al., 2020; Wang et al., 2014). Lastly, the pro-degenerative effects of inflammation in the setting of neonatal HI are not fully alleviated by therapeutic hypothermia (Martinello et al., 2022; Osredkar et al., 2014). Therefore, it will also be important to establish in future studies whether hCG confers added therapeutic benefit in a similar preclinical neonatal injury model of combined cooling with hCG.

## Ethical Publication Statement

We confirm that we have read the Journal’s position on issues involved in ethical publication and affirm that this report is consistent with those guidelines.

## Acknowledgements

This work was supported by the National Institutes of Health R01 NS112234 and the Hope Center for Neurological Disorders at Washington University in St. Louis.

